# Streamline unsupervised machine learning to survey and graph indel-based haplotypes from pan-genomes

**DOI:** 10.1101/2023.02.11.527743

**Authors:** Bosen Zhang, Haiyan Huang, Laura E. Tibbs-Cortes, Adam Vanous, Zhiwu Zhang, Karen Sanguinet, Kimberly A. Garland-Campbell, Jianming Yu, Xianran Li

**Author notes:** Correspondence. Xianran Li.

## Abstract

Identification and visualization of large insertion and deletion (indel) polymorphisms, which contribute significantly to natural phenotypic variation, are challenge from a pan-genome. Here, through streamlining two unsupervised machine learning algorithms, we developed a BRIDGEcereal webapp for surveying and graphing indel-based haplotypes for genes of interest from publicly accessible pangenomes. Over hundreds of assemblies from five major cereals were compiled. We demonstrated the potential of BRIDGEcereal in exploring natural variation with wheat candidate genes within QTLs and GWAS intervals. BRIDGEcereal is available from https://bridgecereal.scinet.usda.gov.

## Main

Pan-genomes with high quality *de novo* assemblies are shifting the paradigm of biology research in genome evolution, speciation, and function annotation (Shi et al., 2022). Arrays of new bioinformatic tools, ranging from data storage, annotation, to polymorphism identification and visualization, have been developed to capitalize pan-genome resources. Large insertion and deletion (indel) polymorphisms, contributing to phenotypic variations thought altering gene structure or expression (Chen et al., 2021), are class of structural variants to be catalogued from pan-genomes. However, for specific genes, surveying and graphing large indels across assemblies are challenge and painstaking tasks (Mahmoud et al., 2019). Here, we constructed an interactive webapp BRIDGEcereal (https://bridgecereal.scinet.usda.gov/) to expedite this process through streamlining unsupervised learning. With multiple wheat genes, we further demonstrated that mining pan-genome through BRIDGEcereal could accelerate gene discovery and characterization.

We devised two unsupervised machine learning algorithms to streamline the process (Figure 1A). The first algorithm, Clustering HSPs for Ortholog Identification via Coordinates and Equivalence (CHOICE, Figure 1B), identifies and extracts the segment harboring the ortholog from each assembly. The segments are then subjected an All-vs-All comparison to survey potential large indels. The second algorithm, Clustering via Large-Indel Permuted Slopes (CLIPS, Figure 1C and Figure S1), groups assemblies sharing the same set of indels to graph a concise haplotype plot to visualize potential large indels, their impacts on the gene, and relationships among haplotypes. For indels outside of genes, because of unknown sizes and locations, multiple iterations (an iteration takes < 20 seconds) are needed for each gene to obtain the optimal haplotype graph by probing different up- and down-stream search boundaries and the order of haplotypes (Figure S2). Through the interactive user-interface of BRIDGEcereal, these parameters can be instantly adjusted based on the visual inspection.

**Figure 1.**
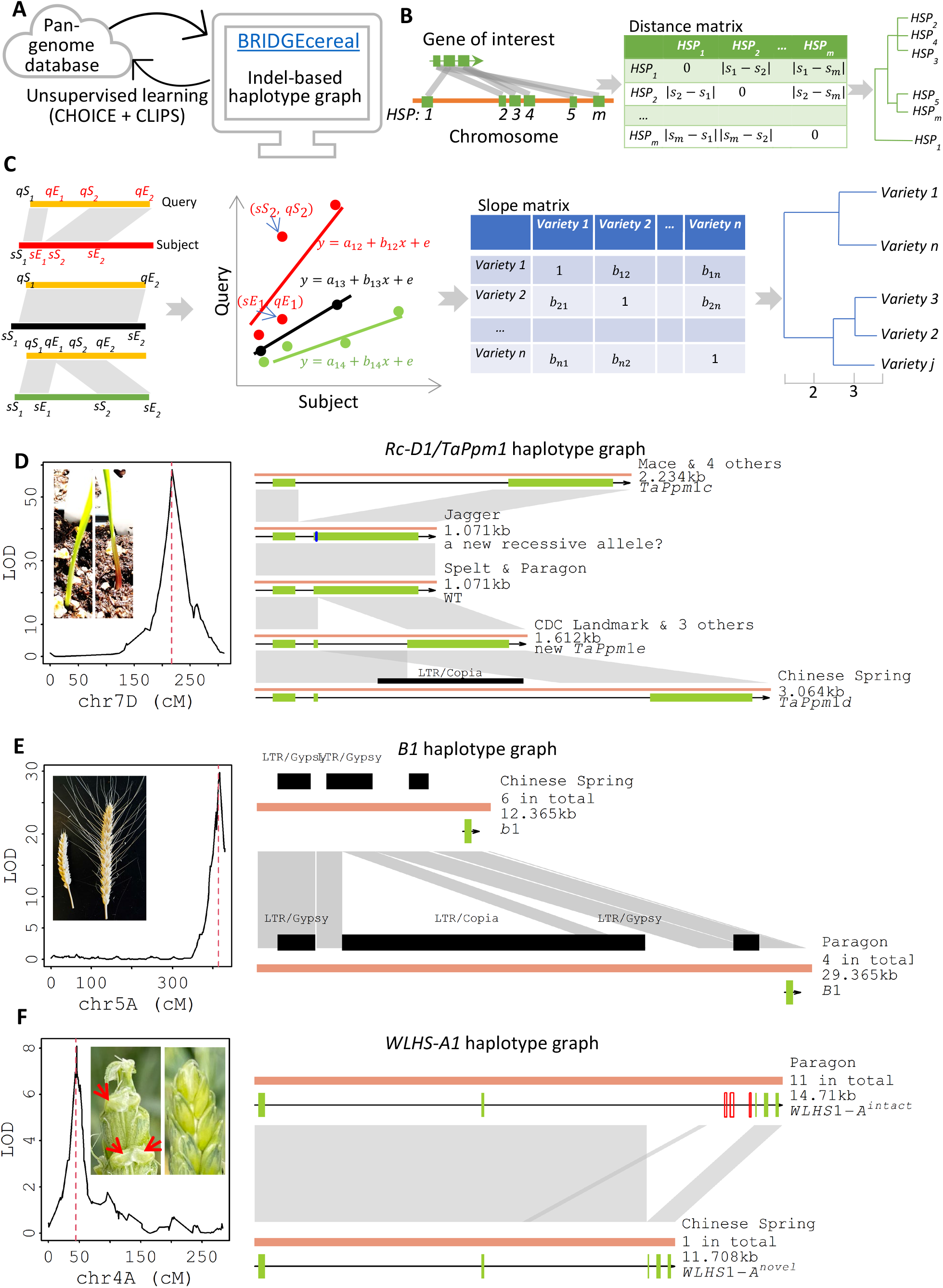
BRIDGEcereal for uncovering large indels and graphing haplotypes for genes of interest. **(A)**. Two unsupervised learning algorithms (CHIOICE and CLIPS) streamline the backend of user-friendly webapp. **(B)**. Depiction of CHOICE. A gene typically has multiple High-scoring Segment Pairs (HSPs, grey polygons). HSPs are clustered with the distance matrix compiled from start coordinates (*s*_*m*_) of the subjects. For each group of clustered HSP, the total HSP length and the mean percentage of identical matches are jointly used to determine the ortholog. In this illustrative case, the ortholog is within the region with HSP2-4. **(C)**. Depiction of CLIPS. Concatenated coordinates of HSPs between two varieties are fit into a linear regression model to estimate the slope. Slope for a pair of varieties without large indels (black) is close to 1 and deviates from 1 with large indels (red or green). The phylogenic tree based on the slope matrix serves as a decision guide to determine the number of haplotypes. **(D)**. Haplotype graph for *Rc-D1/TaPpm1* underlying the coleoptile color QTL. The blue vertical bar marks the nonsynonymous SNP. (**E**). Haplotype graph showing a potential causal large indels regulating the expression of *B1* underlying the major awn type QTL on chromosome 5A. (**F**). Haplotypes graph of *WLHS1-A* located in the *Hd* interval. Three exons (red open boxes) were deleted in Chinese Spring. In D-F, the dashed red vertical line marks the projected position of candidate gene in the QTL interval associated with the corresponding phenotypic variations. Brown lines denote extracted segments harboring the ortholog, which is indicated by green boxes; HSPs are indicated by the grey polygons, while indels by white. Black boxes denote sequences similar to transposons. Arrow indicates gene orientation.

We compiled 108 assemblies of wheat, barley, maize, sorghum, and rice from literature (Table S1). Of these five cereal crops, wheat has the most complex genome: hexaploid with 12-Gb repetitive sequences out of 15-Gb genome. With wheat genes within three QTL intervals and one GWAS signal, we showed that the surveying and graphing large indels from pan-genomes could narrow down candidate genes and generate testable hypotheses.

All three QTLs were detected with the Paragon × Chinese Spring recombinant inbred line population (269 individuals genotyped with 9,434 SNPs). For each measured trait (coleoptile color, awn type, and hooded, Table S2), we identified one major QTL with *prior* genetic information ranged from the identified gene with sound casual sites, the identified gene with unknown causal sites, to an unidentified gene. Potential large indels were surveyed for genes within the QTL interval among 11 assemblies of diverse wheat varieties via BRIDGEcereal (Walkowiak et al., 2020). With uncovered haplotypes, scaffolds (*n* = 1,254,002) of Paragon were searched to reveal its specific haplotype.

We detected one strong QTL on chromosome 7D for coleoptile color (Figure 1D and Figure S3A). The candidate gene was *Rc-D1*/*TaPpm1* (TraesCS7D02G166500) with three reported mutations within the gene (Jiang et al., 2018). We searched boundary was 100-bp upstream and 100-bp downstream. Among 11 varieties, four haplotypes attributed to three indels within the gene were uncovered (Figure 1D and Table S3). Paragon and Spelt (red coleoptile) had the same haplotype as the wild type. No indel was detected in Jagger with green coleoptile, implying that the missense mutation (G50S, Figure S4) was likely a new recessive allele. Five varieties had the *TaPpm1c* allele with a 1.163-kb insertion in the intron. Chinese Spring carries the *TaPpm1d* allele with a 1.993-kb insertion in the second exon. Three varieties shared another new recessive allele (*TaPpm1e*) of a 0.541-kb insertion in the second exon. The evolutionary relationship between *TaPpm1d* and *TaPpm1e* demonstrated the importance of user inputs in determining the optimal depiction of uncovered indels (Figure S5).

Multiple QTLs were detected for awn presence or absence. The strongest QTL on chromosome 5A corresponded to *Tipped1* (*B1*, Figure 1E and Figure S3B) with Paragon carrying the dominant allele suppressing awn development. TraesCS5A02G542800 was identified as the underlying gene, but the consensus was that an undetermined causal polymorphism (outside of the gene body) altered the expression pattern (DeWitt et al., 2020; Huang et al., 2020). We searched potential large indels within the boundary from 30-kb upstream to 100-bp downstream around this gene. The assembled chromosome 5A of Julius and CDC Stanley did not harbor the ortholog. The remaining nine assemblies could be clustered into two haplotypes due to a 17-kb segment containing two indels (14-kb and 3-kb) 6-kb upstream (Figure 1E). The ends of two scaffolds containing part of the 17-kb insertion supported that Paragon shared the same haplotype as ArinaLrFor (Figure S6). This 17-kb segment potentially encodes regulatory elements promoting the expression of *B1*, which is a hypothesis testable by gene editing technologies.

We identified two significant QTL for hooded phenotype (Figure S3C). The QTL on chromosome 4A corresponded to *Hooded* (*Hd*, Figure 1F), which also had a significant epistatic interaction with *B1* for awn type (Figure S3D). The underlying gene for *Hd* remains unknown. In the QTL interval, a homolog (*TaDL*, TraesCS4A02G058800) of rice *dropping leaf* was suggested as the candidate gene (Yoshioka et al., 2017). CRISPR-induced null alleles of *TaDL* suppressed awn development in wheat (Du et al., 2021), further supported this hypothesis. Within the 100-kb segment surrounding *TaDL*, a 12-kb insertion 25-kb downstream of the gene was detected in LongReach Lancer. However, we found that Paragon shared the same haplotype with Chinese Spring (Figure S7 and Table S3). Meanwhile, for TraesCS4A02G058900 (*WLHS1-A*, an ortholog of *OsMADS1*), located 1.5-Mb downstream of *TaDL*, we found that a 2.995-kb deletion in Chinese Spring removed 3 exons, while Paragon and all other varieties shared another haplotype without the deletion (Figure 1F and Table S3). We further aligned RNAseq reads sampled from NongDa3338, Azhurnaya, and Chinese Spring to Spelt and verified the gene structure, its high expression in awn and lemma, and the expression dynamics during spike development (Figure S8). The previous study characterized these two alleles as *WLHS1-A*^*nove*^ and *WLHS1-A*^*intact*^ but didn’t report any associated phenotypic variation (Shitsukawa et al., 2007). Results from BRIDGEcereal mining pan-genomes, combined with phenotypes of *OsMADS1* mutants (Jeon et al., 2000), suggested that *WLHS1-A*, rather than *TaDL*, more likely corresponded to *Hd*.

A recent GWAS identified a significant signal for wheat plant height on chromosome 2A with exome-capture SNPs. Within the 3Mb surveyed region, *TaARF12* (TraesCS2A02G547800) was reported as the candidate. The function of *TaARF12* in regulating plant height was supported by results from RNAi experiments, but causal polymorphisms was not investigated (Li et al., 2022). Via BRIDGEcereal, we identified a *Copia* transposon (6-kb) insertion upstream of *TaARF12* (Figure S9 and Table S3). A follow-up experiment to test whether this is a causal polymorphism would be evaluating the gene expression pattern difference among varieties segregating this transposon.

In summary, by devising two unsupervised learning algorithms, we constructed the BRIDGEcereal webapp as a one-stop gateway to efficiently mine publicly accessible cereal pan-genomes to prioritize candidate genes within QTL/GWAS intervals and explore full spectrum of polymorphisms.

## Acknowledgements

The authors thank the Germplasm Resource Unit at John Innes Centre, UK (Grant number BBS/E/J/000PR8000) for providing the Paragon × Chinese Spring RIL population and sharing the genotype and genetic map information, and thank USDA-ARS SCINet for computing resource and the collaboration of the USDA-ARS-Partnerships for Data Innovations (PDI, https://pdi.scinet.usda.gov/), which provided data stewardship solutions to enable secure data management, storage and sharing.

## Author contributions

X.L. designed the research. B.Z., H.Y., L.E.T, and X.L. conducted the research. Z.Z., K.S., A.V., K.A.G. and J.Y. contributed materials and analysis tools and interpreted results. X.L. and B.Z. wrote the manuscript with inputs from all authors.

## Data and materials availability

All data are available in the main text, the supplementary materials, or https://bridgecereal.scinet.usda.gov.

## Supplementary Methods

### Pan-genomes for five cereals

FASTA files of assemblies at the pseudochromosome level were downloaded from URLs listed in the original publications and split into individual chromosome files (Table S1). After the blast database for each chromosome was built through *makeblastdb*, the FASTA file was compressed into the bgzf format by *bgzip* then indexed with *faidx* (Danecek et al., 2021). For privacy concerns, the uploaded chromosome sequence must use an identifier of either ‘Parent 1’ or ‘Parent 2’, for the FASTA file to disassociate the variety identity.

### CHOICE (Clustering HSP for Ortholog Identification via Coordinates and Equivalency)

A transcript sequence, either retrieved from the assemble (via a gene model ID) or submitted via a FASTA sequence, typically has multiple high-scoring-pairs (HSPs) within each assembly because of multiple exons and partial regions matching to other sequence on the same chromosome. To determine the ortholog for the submitted gene, a distance matrix based on the start coordinates, the 9^th^ column (sStart) of the *Blastn* output with -outfmt 6, of all HSPs will be constructed to build a clustering tree of the HSPs. The HSPs are then split into groups based on the adjustable distance cutoff (the default is 20-kb). For each group, the ratio of the total length of HSPs to the transcript length of the submitted gene will be calculated. For the groups with the ratio within the user-defined range (the default is 0.75 to 1.25), the group with the highest average similarity (the 3^rd^ column of the *Blastn* output) is selected as the ortholog. All the extracted segments containing the ortholog are adjusted to the same orientation.

### CLIPS (Clustering via Large Indel Permuted Slopes)

Coordinates of HSPs between each pair of extracted segments are fit into a linear regression model *y* = *a* + *bx* + *e* to estimate the slope *b*, where *y* is the concatenation of the 7^th^ (qStart) and 8^th^ (qEnd) column from the *Blastn* report (-outfmt 6), and *x* is the concatenation of the 9^th^ (sStart) and 10^th^ (sEnd) column. The estimated slope is close to 1 for a pair of varieties without large indels, greater than 1 for a large insertion in the query, and less than 1 with a large deletion in the query. HSPs attributed to repetitive sequences were removed for the slope estimation (Figure S10). This slope matrix is used to build a clustering tree of all varieties. The varieties sharing the same set of indels will be clustered together. A user selected cutoff (via clicking on the tree plot) is used to define the haplotypes. One variety is randomly selected as the representative for graphing the haplotypes.

### Phenotype and QTL mapping

Seeds of the Paragon × Chinese Spring RIL population were received from the Germplasm Resource Unit at John Innes Centre (Gardiner et al., 2019). The genotype (via the 35K Axiom SNP array) and genetic map information were retrieved from CereaslDB.com. The entire population was grown in a greenhouse. Coleoptile color was visually scored after seedling emerged from soil as red for 1 and green for 0. Awn type was visually scored as presence (1) or absence (0) at maturity. The hooded phenotype was scored following previous report as 1 for presence of a membranous structure or broadening awn base and 0 for neither (Yoshioka et al., 2017). R/qtl was used for identifying QTL and epistatic interaction (Broman et al., 2003). The physical location of the QTL interval was determined by searching the corresponding Axiom SNP probe sequence again the Chinese Spring reference v1.0. RNAseq reads sampled from NongDa3338 (Chen et al., 2022), Azhurnaya and Chinese Spring (Ramirez-Gonzalez et al., 2018) were downloaded from GenBank SRA and mapped to the Spelt genome by HISAT2 v2.2.1 (Kim et al., 2019). Sashimi plot was generated by IGV v2.12.3 (Robinson et al., 2011). The TPM value of *WLHS-A1* across developmental stages were tabulated by featureCounts (Liao et al., 2014).

---

**Table S1.** Assemblies (at the pseudochromosome level) included in BRIDGEcereal.

| Cereal | Variety | Reference |
| --- | --- | --- |
| Wheat<br>(11) | ArinaLrFor; CDC Landmark; CDC Stanley; Chinese Spring; Jagger; Julius; LongReach Lancer; Mace; Norin 61; Spelt; SY Mattis | (Walkowiak et al., 2020) |
| Barley<br>(20) | Akashinriki; B1K_04_12; Barke; Golden Promise; Hockett; HOR 10350; HOR 13821; HOR 13942; HOR 21599; HOR 3081; HOR 3365; HOR 7552; HOR 8148; HOR 9043; Igri; Morex; OUN333; RGT Planet; ZDM01467; ZDM02064 | (Jayakodi et al., 2020) |
| Maize<br>(26) | B73; B97; CML103; CML228; CML247; CML277; CML322; CML333; CML52; CML69; Hp301; Il14H; Ki11; Ki3; Ky21; M162W; M37W; Mo17; Mo18W; MS71; NC350; NC358; Oh43; Oh7B; P39; Tx303; Tzi8 | (Hufford et al., 2021) |
| Sorghum<br>(18) | 353; AusTRCF317961; BTx430; BTx623; BTx642; IS12661; IS19953; IS3614-3; IS8525; IS929; Ji2731; PI525695; PI532566; PI536008; R931945-2-2; Rio; S369-1; SC187 | (Tao et al., 2021) |
| Rice<br>(33) | 2428; 9311; Basmati1; CG14; CN1; D62; DG; DHX2; FH838; FS32; G46; G630; G8; II32; IR64; J4155; Kosh; KY131; Lemont; LJ; N22; NamRoo; NIP; R498; R527; S548; TM; TUMBA; WSSM; Y3551; Y58S; YX1; ZH11 | (Qin et al., 2021) |

**Table S2.** Observed phenotypes for the wheat Paragon × Chinese Spring RIL population (in Excel).

**Table S3.** Haplotypes of candidate genes among wheat assemblies. Note: The Paragon haplotype was determined by an extra search process because its genome assembly is at the scaffold level. For *TaDL* and *TaARF12* without the genetic context, ‘-’ is used for one of haplotypes without insertion.

| Gene | Varieties sharing the same haplotype | Contributing polymorphism |
| --- | --- | --- |
| ***Rc-D1/TaPpm1*** | Spelt; **<u>Paragon</u>** | Wild type |
|  | Jagger | G50S (a new recessive allele?) |
|  | Mace; SY Mattis; ArinaLrFor; Julius; LongReach Lancer | A 1.163-kb insertion (*TaPpm1c*) |
|  | **<u>Chinese Spring</u>** | A 1.993-kb insertion (*TaPpm1d*) |
|  | CDC Landmark; CDC Stanley; Norin 61 | A 0.541-kb insertion (new *TaPpm1e*) |
| ***B1*** | **<u>Paragon</u>**; ArinaLrFor; SY Mattis; Spelt | *B1* |
|  | **<u>Chinese Spring</u>**; Jagger; Norin 61; Mace; LongReach Lancer; CDC Landmark | A 17-kb deletion at 6-kb upstream (*b1*) |
|  | Julius; CDC Stanley | NA |
| ***TaDL*** | **<u>Chinese Spring</u>**; CDC Landmark; ArinaLrFor; CDC Stanley; Jagger; Julius; Norin 61; Mace; Spelt; SY Mattis; **<u>Paragon</u>** | - |
|  | LongReach Lancer | A 13.5-kb insertion at 25-kb downstream |
| ***WLHS1-A*** | **<u>Paragon</u>**; LongReach Lancer; CDC Landmark; ArinaLrFor; CDC Stanley; Jagger; Julius; Norin 61; Mace; Spelt; SY Mattis; | Wild type (*WLHS1-A<sup>intact</sup>*) |
|  | **<u>Chinese Spring</u>** | A 2.995-kb deletion removed 3 exons (*WLHS1-A<sup>novel</sup>*) |
| ***TaARF12*** | Spelt; SY Mattis | - |
|  | CDC Landmark; ArinaLrFor; Chinese Spring; CDC Stanley; Jagger; Julius; LongReach Lancer; Norin 61; Mace | A 6-kb insertion at 4-kb upstream. |

**Figure S1.**
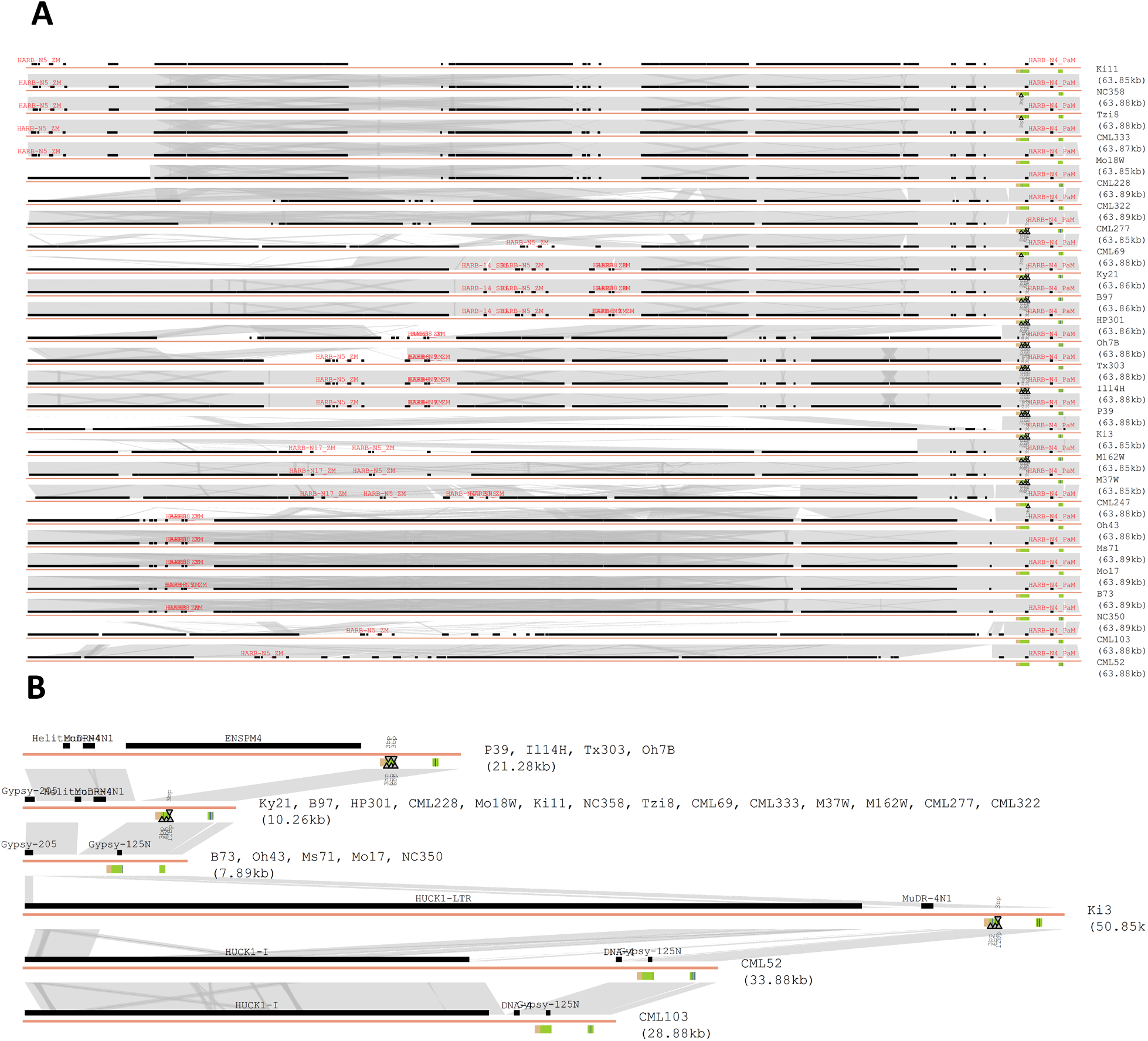
CLIPS for grouping varieties sharing the same indels. (**A**) The indiscernible representation of indels in the upstream region (60-kb) of *ZmCCT9* among 26 maize inbreds before clustering into haplotypes. (**B**) A legible representation of indels of *ZmCCT9* after clustering into 6 haplotypes.

**Figure S2.**
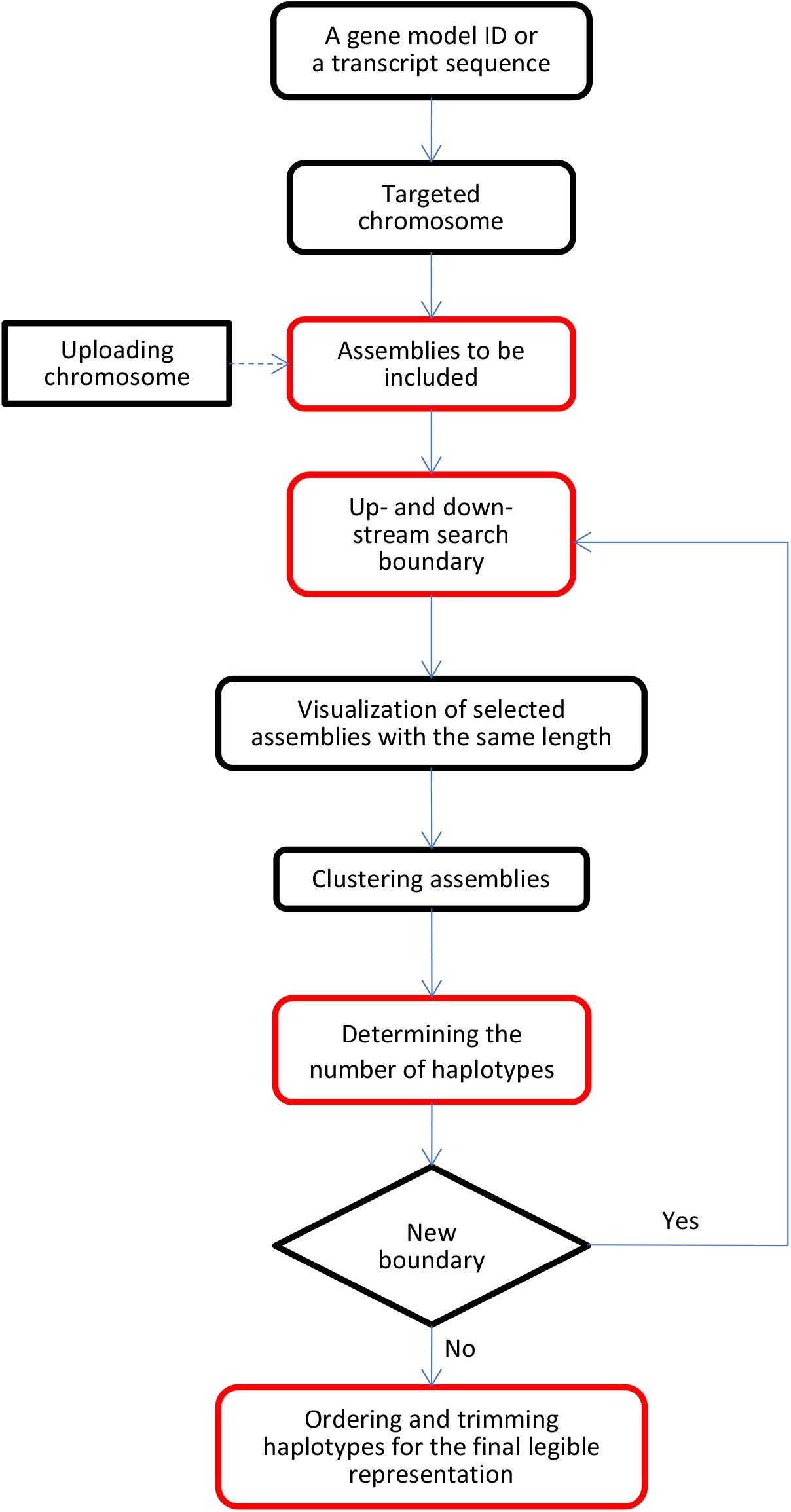
Streamlined workflow for the BRIDGEcereal webapp. Red outline denotes steps with parameter selected based on user’s discretion.

**Figure S3.**
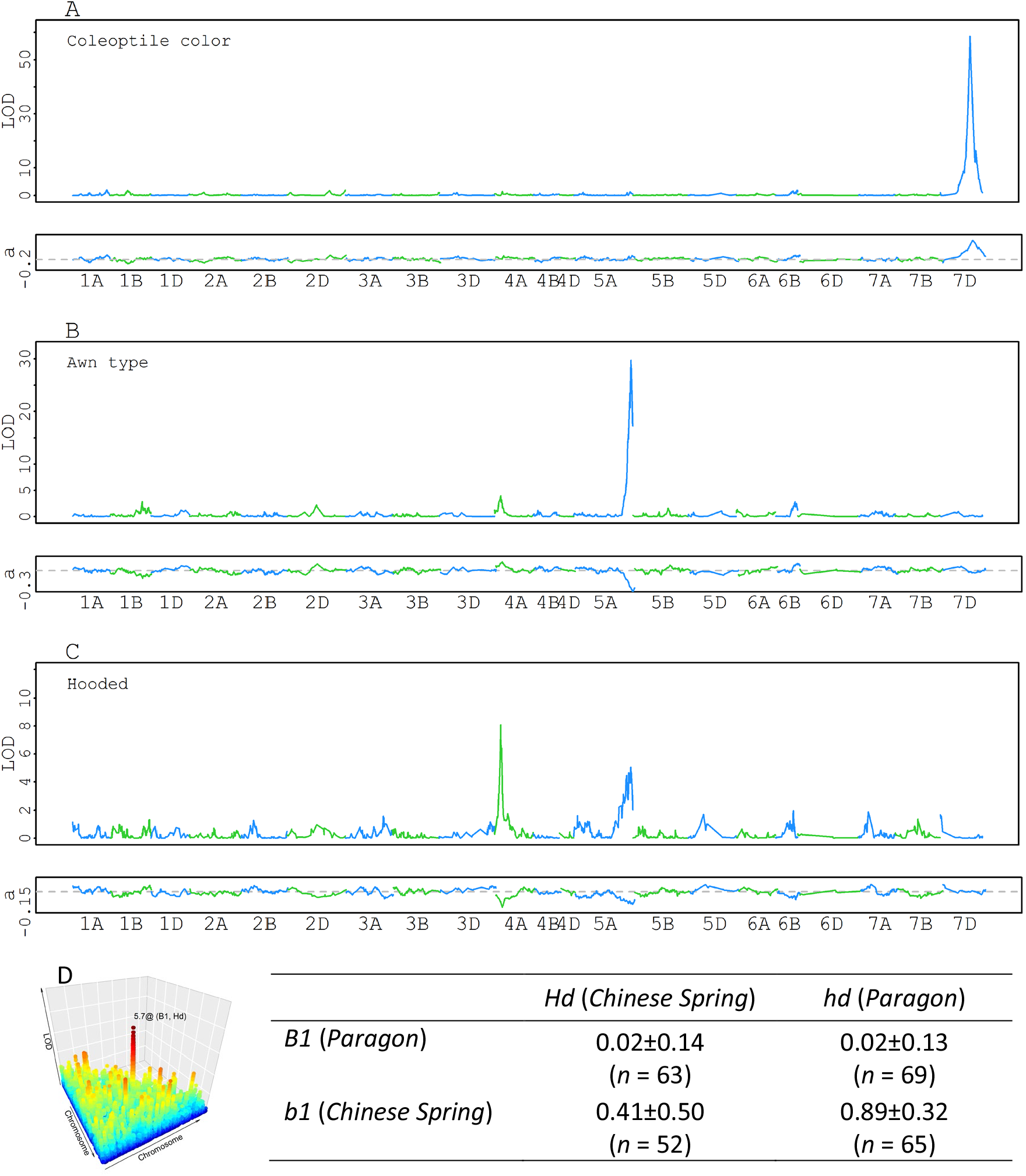
QTLs identified with the Paragon × Chinese Spring RIL population. (**A**) The major QTL on chromosome 7D for coleoptile color (0 for green and 1 for red) corresponded to *Rc-D1*. (**B**) Multiple QTL for awn type (0 for absence and 1 for presence). The major QTL on chromosome 5A corresponded to *B1*. Two minor QTLs on chromosome 4A and 6B corresponded to *Hooded* (*Hd*) and *Tipped 2* (*B2*), respectively. Literature indicated that dominant alleles of three genes inhibit awn development. QTL effects indicated that Chinese Spring carries *Hdb1B2*, while Paragon carries *hdB1b2*, which agreed with the observation that both are an awnletted type with short awn. (**C**). *Hd* (chromosome 4A) and *B1* were two significant QTLs for the hooded phenotype (0 for non-hooded and 1 for hooded). (**D**). Epistasis interaction between *B1* and *Hd* for the awn type. Each cell shows the mean and standard deviation for the numerical score for the awn type. For each SNP, the Chinese Spring allele is coded as 0 while the Paragon allele as 1. The dashed gray horizontal line marks the 0 effect.

**Figure S4.**
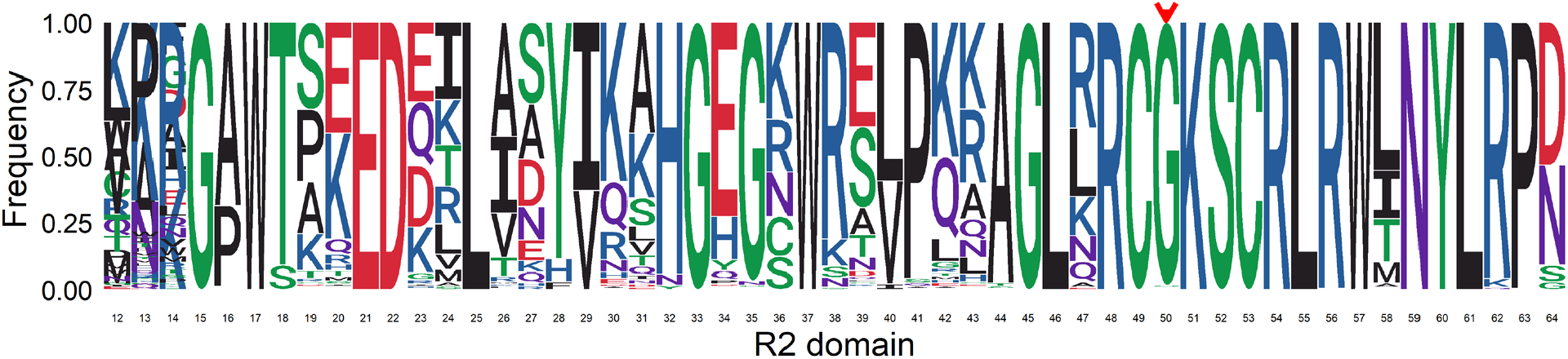
Jagger had a G50S missense (red arrowhead) mutation within the highly conserved R2 domain of *Rc-D1*/*TaPpm1*. The frequency of each amino acid is calculated from 247 proteins. This missense mutation is the only polymorphism detected between Jagger (green coleoptile) and the wild type, implying it is a potential new recessive allele.

**Figure S5.**
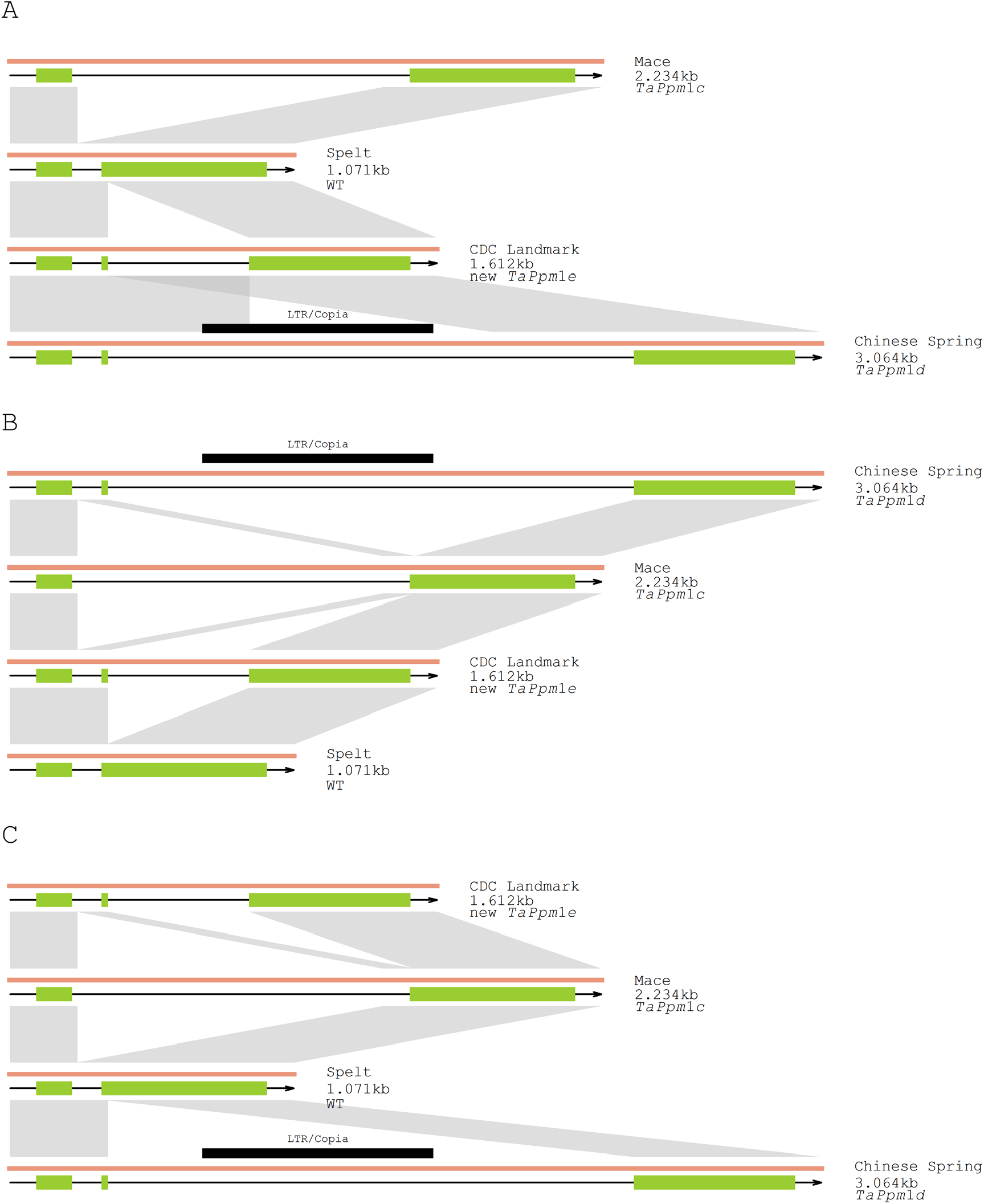
The interactive GUI incorporates the users’ input to determine how the haplotypes are presented. Different orders of haplotypes may lead to different perception and interpretations. For example, the evolutionary relationship between *TaPpm1d* and the new allele *TaPpm1e* could be clearly revealed in **A**, but not in **B** and **C**, two random orders out of 24 possibilities.

**Figure S6.**
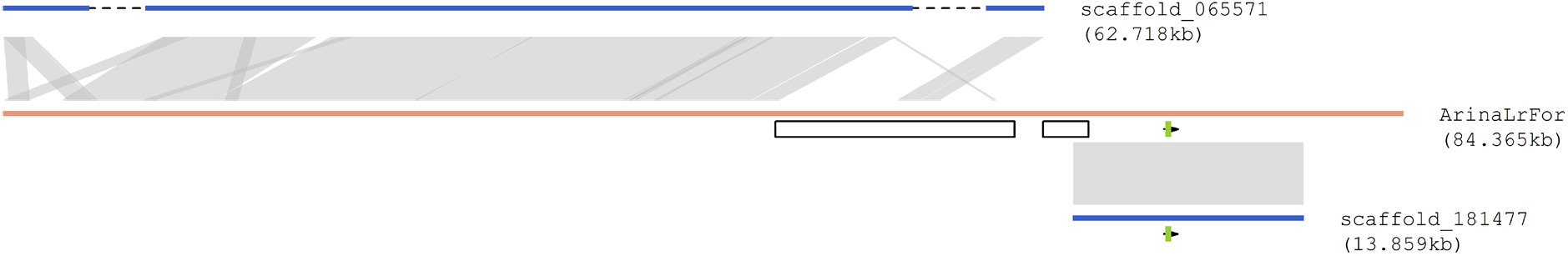
Paragon harbors the 17-kb segment in the upstream of *B1*. The ends of two scaffolds (blue) have partial sequences of the 17-kb segments assembled in ArinaLrFor. The green box denotes the zinc finger gene underlying *B1*. The two open black boxes are the 14-kb and 3-kb segments deleted in Chinese Spring (Figure 2B), respectively. Both scaffolds (in full length) are reversed and complemented for the plotting purposes. The dashed black lines in scaffold_065571 are large sequence gaps.

**Figure S7.**
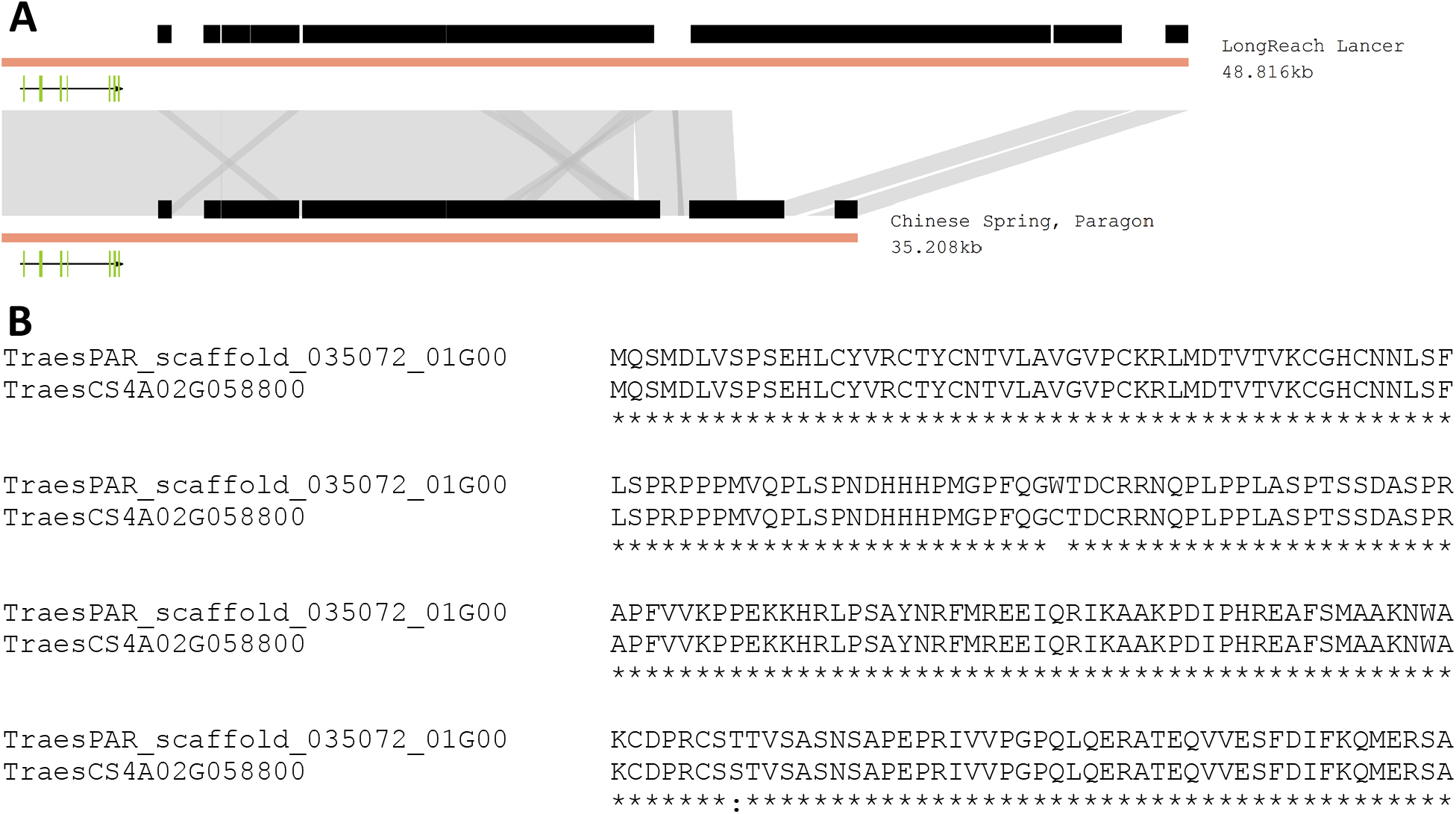
*TaDL* is unlikely the underlying gene of *Hd*. **(A**). A 13.5-kb insertion at the downstream 24-kb was detected in LongReach Lancer, but Chinese Spring and Paragon shared the same haplotype. **(B)**. Alignment for the TaDL protein sequences between Paragon and Chinese Spring.

**Figure S8.**
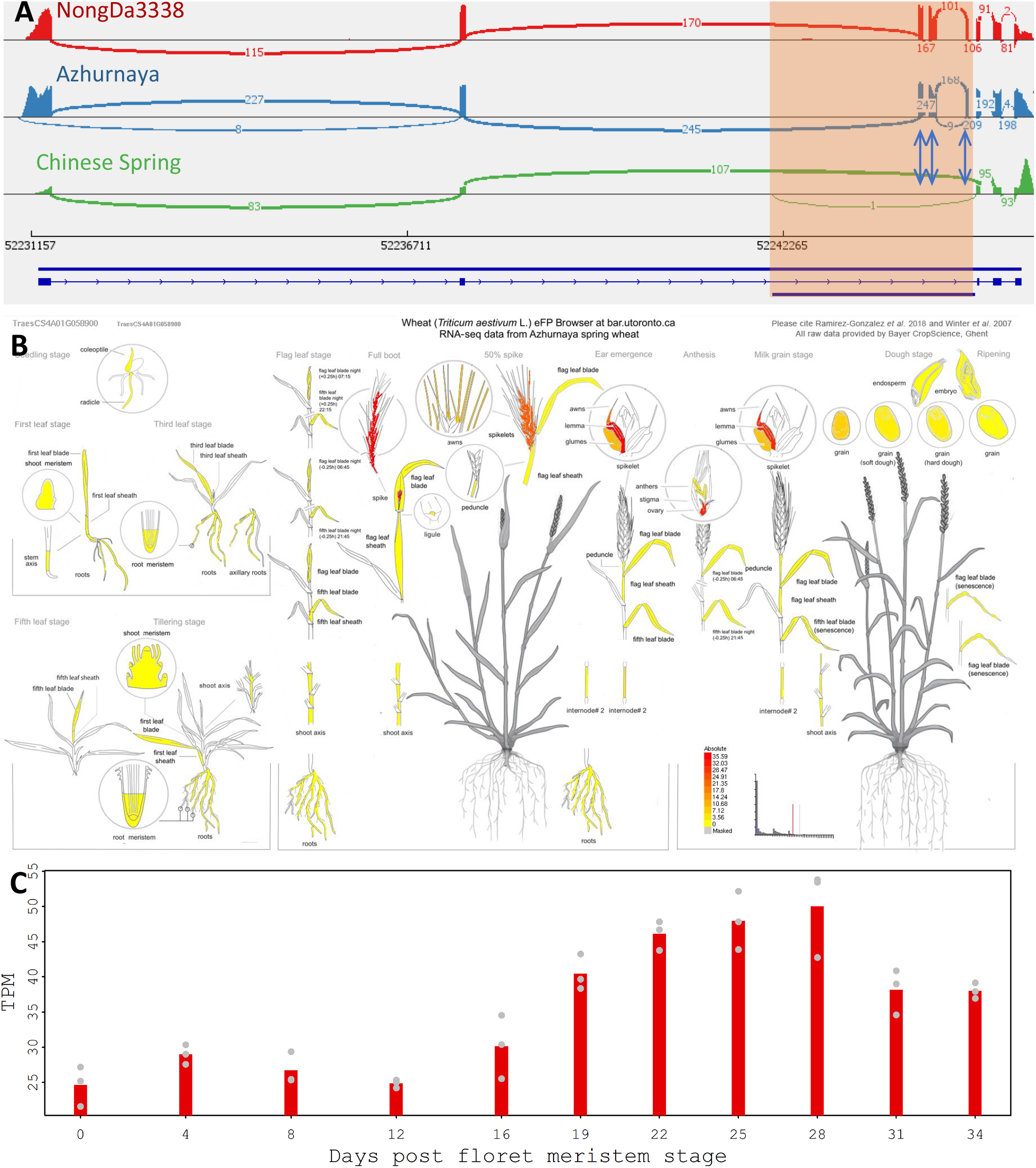
Structure and expression pattern of *WLHS1-A*^*intact*^. (**A**). RNAseq reads from NongDa338 (SRR19046544), Azhurnaya (ERR2409914), and Chinese Spring (SRR6802609 and SRR6802614) mapped to the Spelt genome. The 2.996-kb deleted segment in Chinese Spring (*WLHS1-A*^*intact*^) includes three exons. The depth was not normalized by total mapped reads in these three tracks. (**B**). *WLHS1-A*^*intact*^ is highly specifically expressed in Awn related tissues (Azhurnaya reads mapped to the Chinese Spring genome, https://bar.utoronto.ca/efp_wheat/cgi-bin/efpWeb.cgi). (C). *WLHS1-A*^*intact*^ expression dynamics during spikelet development in NongDa3338. TPM: Transcripts Per Kilobase Million.

**Figure S9.**
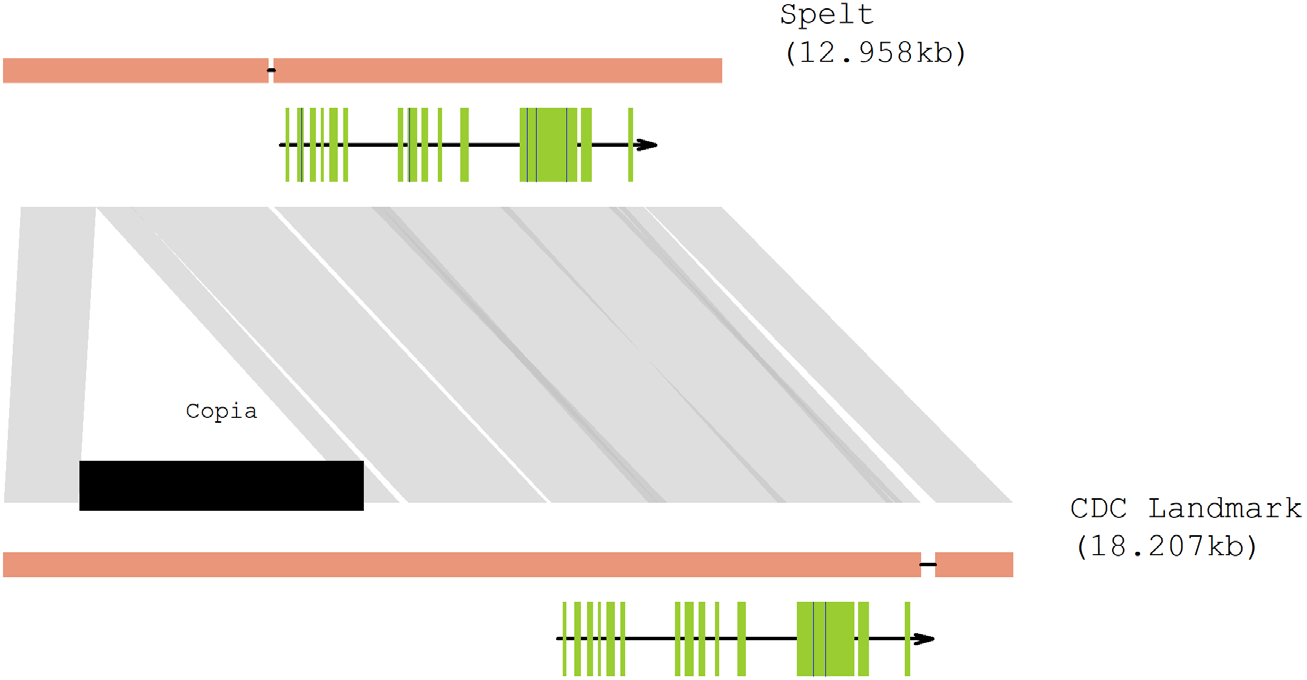
A 6-kb insertion 4-kb upstream of *TaARF12* might be the causal site. *TaARF12* was detected in a GWAS for plant height with exome capture SNPs, but the causal polymorphism has not been investigated.

**Figure S10.**
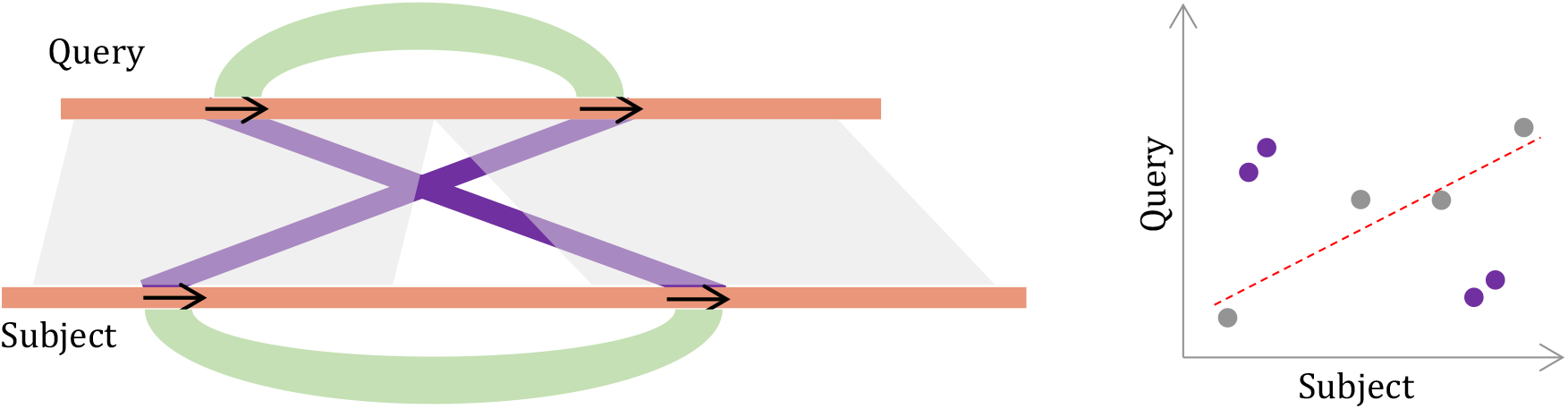
HSPs (purple polygons) attributed to repetitive sequences (black arrows) were excluded before the slope estimation. These HSPs have a substantial weight in the slope estimation and may diminish the slope changed introduced by indels. These HSPs were identified by cross-referencing to HSPs (green curves) from comparing each segment against itself, which are included in the second-round *Blastn*.

## References

Chen, Q., Li, W., Tan, L., and Tian, F. (2021). Harnessing knowledge from maize and rice domestication for new crop breeding. Mol Plant 14:9–26. 10.1016/j.molp.2020.12.006.

DeWitt, N., Guedira, M., Lauer, E., Sarinelli, M., Tyagi, P., Fu, D., Hao, Q., Murphy, J.P., Marshall, D., Akhunova, A., et al. (2020). Sequence-based mapping identifies a candidate transcription repressor underlying awn suppression at the B1 locus in wheat. New Phytol 225:326–339. 10.1111/nph.16152.

Du, D., Zhang, D., Yuan, J., Feng, M., Li, Z., Wang, Z., Zhang, Z., Li, X., Ke, W., Li, R., et al. (2021). FRIZZYPANICLE defines a regulatory hub for simultaneously controlling spikelet formation and awn elongation in bread wheat. New Phytol 231:814–833. 10.1111/nph.17388.

Huang, D., Zheng, Q., Melchkart, T., Bekkaoui, Y., Konkin, D.J.F., Kagale, S., Martucci, M., You, F.M., Clarke, M., Adamski, N.M., et al. (2020). Dominant inhibition of awn development by a putative zinc-finger transcriptional repressor expressed at the B1 locus in wheat. New Phytol 225:340–355. 10.1111/nph.16154.

Jeon, J.S., Jang, S., Lee, S., Nam, J., Kim, C., Lee, S.H., Chung, Y.Y., Kim, S.R., Lee, Y.H., Cho, Y.G., et al. (2000). leafy hull sterile1 is a homeotic mutation in a rice MADS box gene affecting rice flower development. Plant Cell 12:871–884. 10.1105/tpc.12.6.871.

Jiang, W., Liu, T., Nan, W., Jeewani, D.C., Niu, Y., Li, C., Wang, Y., Shi, X., Wang, C., Wang, J., et al. (2018). Two transcription factors TaPpm1 and TaPpb1 co-regulate anthocyanin biosynthesis in purple pericarps of wheat. J Exp Bot 69:2555–2567. 10.1093/jxb/ery101.

Li, A., Hao, C., Wang, Z., Geng, S., Jia, M., Wang, F., Han, X., Kong, X., Yin, L., Tao, S., et al. (2022). Wheat breeding history reveals synergistic selection of pleiotropic genomic sites for plant architecture and grain yield. Mol Plant 15:504–519. 10.1016/j.molp.2022.01.004.

Mahmoud, M., Gobet, N., Cruz-Davalos, D.I., Mounier, N., Dessimoz, C., and Sedlazeck, F.J. (2019). Structural variant calling: the long and the short of it. Genome Biol 20:246. 10.1186/s13059-019-1828-7.

Shi, J., Tian, Z., Lai, J., and Huang, X. (2022). Plant pan-genomics and its applications. Mol Plant 10.1016/j.molp.2022.12.009.

Shitsukawa, N., Tahira, C., Kassai, K., Hirabayashi, C., Shimizu, T., Takumi, S., Mochida, K., Kawaura, K., Ogihara, Y., and Murai, K. (2007). Genetic and epigenetic alteration among three homoeologous genes of a class E MADS box gene in hexaploid wheat. Plant Cell 19:1723–1737. 10.1105/tpc.107.051813.

Walkowiak, S., Gao, L., Monat, C., Haberer, G., Kassa, M.T., Brinton, J., Ramirez-Gonzalez, R.H., Kolodziej, M.C., Delorean, E., Thambugala, D., et al. (2020). Multiple wheat genomes reveal global variation in modern breeding. Nature 588:277–283. 10.1038/s41586-020-2961-x.

Yoshioka, M., Iehisa, J.C.M., Ohno, R., Kimura, T., Enoki, H., Nishimura, S., Nasuda, S., and Takumi, S. (2017). Three dominant awnless genes in common wheat: Fine mapping, interaction and contribution to diversity in awn shape and length. PLoS One 12:e0176148. 10.1371/journal.pone.0176148.

## References

Broman, K.W., Wu, H., Sen, S., and Churchill, G.A. (2003). R/qtl: QTL mapping in experimental crosses. Bioinformatics 19:889–890. 10.1093/bioinformatics/btg112.

Chen, Y., Guo, Y., Guan, P., Wang, Y., Wang, X., Wang, Z., Qin, Z., Ma, S., Xin, M., Hu, Z., et al. (2022). A wheat integrative regulatory network from large-scale complementary functional datasets enables trait-associated gene discovery for crop improvement. Mol Plant 10.1016/j.molp.2022.12.019.

Danecek, P., Bonfield, J.K., Liddle, J., Marshall, J., Ohan, V., Pollard, M.O., Whitwham, A., Keane, T., McCarthy, S.A., Davies, R.M., et al. (2021). Twelve years of SAMtools and BCFtools. Gigascience 10 10.1093/gigascience/giab008.

Gardiner, L.J., Wingen, L.U., Bailey, P., Joynson, R., Brabbs, T., Wright, J., Higgins, J.D., Hall, N., Griffiths, S., Clavijo, B.J., et al. (2019). Analysis of the recombination landscape of hexaploid bread wheat reveals genes controlling recombination and gene conversion frequency. Genome Biol 20:69. 10.1186/s13059-019-1675-6.

Hufford, M.B., Seetharam, A.S., Woodhouse, M.R., Chougule, K.M., Ou, S., Liu, J., Ricci, W.A., Guo, T., Olson, A., Qiu, Y., et al. (2021). De novo assembly, annotation, and comparative analysis of 26 diverse maize genomes. Science 373:655–662. 10.1126/science.abg5289.

Jayakodi, M., Padmarasu, S., Haberer, G., Bonthala, V.S., Gundlach, H., Monat, C., Lux, T., Kamal, N., Lang, D., Himmelbach, A., et al. (2020). The barley pan-genome reveals the hidden legacy of mutation breeding. Nature 588:284–289. 10.1038/s41586-020-2947-8.

Kim, D., Paggi, J.M., Park, C., Bennett, C., and Salzberg, S.L. (2019). Graph-based genome alignment and genotyping with HISAT2 and HISAT-genotype. Nat Biotechnol 37:907–915. 10.1038/s41587-019-0201-4.

Liao, Y., Smyth, G.K., and Shi, W. (2014). featureCounts: an efficient general purpose program for assigning sequence reads to genomic features. Bioinformatics 30:923–930. 10.1093/bioinformatics/btt656.

Qin, P., Lu, H., Du, H., Wang, H., Chen, W., Chen, Z., He, Q., Ou, S., Zhang, H., Li, X., et al. (2021). Pan-genome analysis of 33 genetically diverse rice accessions reveals hidden genomic variations. Cell 184:3542–3558 e3516. 10.1016/j.cell.2021.04.046.

Ramirez-Gonzalez, R.H., Borrill, P., Lang, D., Harrington, S.A., Brinton, J., Venturini, L., Davey, M., Jacobs, J., van Ex, F., Pasha, A., et al. (2018). The transcriptional landscape of polyploid wheat. Science 361 10.1126/science.aar6089.

Robinson, J.T., Thorvaldsdottir, H., Winckler, W., Guttman, M., Lander, E.S., Getz, G., and Mesirov, J.P. (2011). Integrative genomics viewer. Nat Biotechnol 29:24–26. 10.1038/nbt.1754.

Tao, Y., Luo, H., Xu, J., Cruickshank, A., Zhao, X., Teng, F., Hathorn, A., Wu, X., Liu, Y., Shatte, T., et al. (2021). Extensive variation within the pan-genome of cultivated and wild sorghum. Nat Plants 7:766–773. 10.1038/s41477-021-00925-x.

